# How long is long enough? Decreasing on plant extract lethal effects, over time, affecting *Aedes aegypti* larval mortality

**DOI:** 10.1101/808980

**Authors:** Gilberto Dinis Cozzer, Renan de Souza Rezende, Junir Antônio Lutinski, Walter Antônio Roman Junior, Maria Assunta Busato, Jacir Dal Magro, Daniel Albeny-Simões

## Abstract

The mosquito *Aedes aegypti* has overcome all kinds of human being mosquito control attempts over the last century. Strategies for vector population control resorts to the use of synthetic insecticides, which can lead to problems of intoxication in humans and environmental contamination. We evaluated the effects of *Bacillus thuringiensis* var. *israelensis* (Bti), *Ilex paraguariensis* (mate-herb) and *Ilex theezans* (caúna-herb) extracts against *A. aegypti* larvae mortality. The bioassays were conducted under controlled laboratory conditions of temperature (27±3°C) and photoperiod (12h). Hydroalcoholic *I. theezans* leaves extract displayed better residual effect compared to *I. paraguariensis* fruit aqueous extract. Variation in larval mortality was also observed in the exposure periods (low after a few weeks). Low mortality after a few weeks may mean increased the food for mosquito in a oppose effect over time. The residual effect of *Bti* was observed during the 56 days of the study duration (100% of mortality). The strongest residual effect of *I. theezans* was probably due to the presence of chemical on its leaves, such coumarins, hemolytic saponins and cyanogenic glucosides, absent in *I. paraguariensis*. On the other hand, alternative methods to vector control present risks in a long term scale by reversal of larvicide effect into food resource. Our results contributed to the prospection of natural insecticides and open the possibility for subsequent studies of the use of plant extracts in field situations in a short time scale.

## Introduction

Over the last century the mosquito *Aedes aegypti* Linnaeus, 1762 has overcome all mosquito control attempts by human being. *Aedes* females are well known by their capacity of naturally and/or under laboratory conditions replicate and transmit over 100 kinds of viruses [1]. As an example, in Brazil dengue, chikungunya, zika and most recently mayaro [1] viruses are a real threat to public health [2]. It became necessary the *A. aegypti* population control in order to reduce viruses’ transmission and consequently the epidemic status. Although several chemical and natural products have being extensively used on attempts to reduce adults and immature mosquito population[3,4], appropriated mosquito population reduction is not close to come true. Especially because the gene resistance selectivity due to chemicals and natural products application misuse [5]. Accordingly, the most effective disease prevention method still remains targeting the mosquito population by elimination of mosquito breeding places [6].

A very promising field for mosquito population reduction is focusing the mosquito control strategies on the immature aquatic stages, when the insect is more vulnerable[7,8]. For this purpose, the use of synthetic insecticides is well known as being quite effective causing mosquito larvae mortality [9–13]. However, those chemicals might affect humans causing intoxication and environmental contamination, affecting biodiversity [9,10,12,13]. For the environment, the continuous use of synthetic insecticides may present undesirable effects, such as the long-term residence in the environment, selection of resistant populations and the appearance of new pests [10,12,13]. Regarding to the human health, the presence of such synthetic chemicals on the environment can cause neurological damage, and is associated with a wide range of symptoms, with significant deficits in nervous system function [14].

Alternatively to the use of synthetic chemicals, biological control play an important role on mosquito control[15,16]. The use of bacteria spores as a mosquito larvicidal has been highlighted among the several components that are part of the mosquito integrated management program [15]. In the last decades, the use of inactivated spores of the *Bacillus thuringiensis* var. *israelensis* (Bti) bacteria spread into the mosquito breeding places water, has accomplished desired results, reaching mortality rates above 99%[17,18]. Additionally, during the last few years, plant derived compounds have been extensively used as an alternative method for controlling mosquitoes, not only because it is a new insecticidal agent, but also because it has been described as being environmental friendly [11,19–21]. The use of natural insecticides has some advantages over traditional synthetic products, since natural products are potentially less toxic to the environment. Environmental friendly compounds are less concentrated, have faster degradation and are specific to certain insect groups, resulting in less occupational exposure and less environmental pollution [22].

*Ilex paraguariensis* A. St.-Hil (Aquifoliaceae), known as “*Mate*” is an abundant plant [23], native of South America, twenty meters tall, endowed with dense crown and very branched [23]. *I. paraguariensis* leaves, after processing, are traditionally used to prepare a regional tea known as “*Chimarrão*” in Argentina, Brazil, Paraguay and Uruguay [23]. *I. paraguariensis* is commercially interesting due to caffeine and theobromine presence, both, recognized by displaying a nervous and cardiocirculatory systems stimulant effect [24]. The described pharmacological activities for *I. paraguariensis* leaf extracts include antioxidant, hypolipidemic[25,26] and hypoglycemic effects [27]. Also, *Ilex theezans* Mart. Ex Reissek (Aquifoliaceae), popularly known as cauna, is commonly found at southern Brazil [23]. It is well known due to the physiological characteristics of its leaves as adulterant of *I. paraguariensis* [28]. It is an evergreen tree, early secondary or late secondary species [23], 20 meters long by 70 cm diameter average [28]. Both, *I. paraguariensis* fruit extract and *I. theezans* hydroacoholic leaves extracts have larvicidal effect against *A. aegypti* larvae[9,20].

There are studies showing that *Ilex* spp. leaves and fruit extracts kill *A. aegypti* larvae within a 24h observational time[9,20]. However, there are no studies on the current literature evaluating the effects of time on the bioinsecticides lethal activity. How long is long enough for the bioinsecticide to still keep the mosquito larvae killing activity? [29–31]. In this context, we evaluated the lethal residual effect of *B. thuringiensis* var. *israelensis*, and leaves and fruits extracts of *I. theezans*, and *I. paraguariensis* (respectively), on *A. aegypti* larvae survivorship. We hypothesized that time would positively affect *A. aegypti* larvae survival due to the plant extracts lethal compounds decay, but the mortality caused by by *I. theezans* would be higher compared to *I. paraguariensis* due to the difference in physical and chemical characteristics.

## Materials and Methods

### Animal source

The *A. aegypti* larvae used in this experiment were provided by the Laboratório de Entomologia Ecológica (LABENT-Eco) mosquito colony. A filter paper holding about 300 eggs was placed in a plastic tray (30×20cm) holding 1L of tap dechlorinated water. After hatching the larvae were split among 3 equal size, above described, plastic tray and feed with 2g of fish food. The mosquito larvae were raised for about 4-5 days until reaching 3^rd^ and ^4rd^ instars.

### Plant source and extracts preparation

Fruits and leaves were obtained from native trees located at the Marechal Bormann district (27 ° 19’05 ‘‘S; 52 ° 65’11’’W), Chapecó (SC), in December, 2016. The plant parts were dehydrated at room temperature (± 20°C), pulverized in a knife mill (Cielamb®, CE 430) and stored away from light and moisture. The plant extracts were done according to Busato *et al*. (2015) and Knakiewicz *et al*. (2016). We took a sample of twenty grams of *I. paraguariensis* dehydrated fruits and *I. theezans* leaves. Both samples were extracted by turbolysis using 200ml of distilled and deionized water and a hydroalcoholic solution (90% ethanol; 200 ml) as the solvent, respectively [32]. The extracts were filtered in Büchner, rotavapor concentrated under reduced pressure, lyophilized, weighed, identified and stored in a freezer at −20°C. Hydroalcoholic and aqueous extracts were prepared using *I. theezans* and *I. paraguariensis* leaves and fruits, respectively. We used leaves in a concentration of 1000µg/ml and fruits were diluted to 2000µg/ml. *B. thuringiensis* var. *israelensis* (Bti) strain WG® was used in a concentration of 0.004g/L, lethal dose specified by the manufacturer.

### Experimental microcosms and design

We used 300ml plastic cups holding 100ml of dechlorinated water plus the treatment proposed. In each individual microcosm we added twenty 3^rd^ and 4^th^ instar *A. aegypti* larvae. Each container was covered with a mosquito net held by a rubber elastic band. We tested for the effects of Bti spores, *I. theezans, I. paraguariensis* leaves and fruits, respectively and clean aged water (control) on the *A. aegypti* larval mortality after seven days exposition. Before running the mortality test, each experimental treatment was aged from one to eight weeks. We considered each week as one age block with each treatment replicated six times. By doing this experimental design we assure the independence of each set of treatments. The aged treatments were used to test for larval survival in each experimental week. At the end of the seventh day the larval survival was recorded. Both pupae and emerged adults were considered survivals. The experiment was performed for eight weeks (56 days) and carried out between April and May 2017 at the LABENT-Eco mosquito colony room under controlled conditions of temperature and photoperiod (27±3°C, 12h D:L).

### Statistics

Because both negative (Bti) and positive (tap water) control survivor rate was 0.16% and 100%, respectively, we analyzed the data in both ways, with (complete model) and without (simple model) these two categories. To evaluate differences in percentage of larval mortality (response variables) in simple (*I. paraguariensis* and *I. theezans*) and complete model (only water, Bti spores, *I. paraguariensis* and *I. theezans*), week (1 to 8) and interaction between week and treatment (explicative variables) we used factorial GLM, with binomial correct to quasi-binomial (link= logit, test= Chi-square) distributions for larval mortality (response variable; [33]). All GLMs analyzed were corrected for cases of under-or overdispersion.

Differences among the categorical variables were assessed trough a contrast analysis [33]. In this contrast analysis (orthogonal), the dependent variables of different treatment and weeks were ordered (increasingly) and tested pairwise (with the closest values). Sequentially these dates are adding to the model values with no differences and testing with the next in a steps model simplification (for more see also chapter 9 of Crawley, (2007)). All analyses were performed using R [34].

## Results

The *A. aegypti* larvae mortality was not affected by water age (positive control). We observed no larva deaths independent of the water age used. Oppositely, the Bti lead to zero percent of larvae survivorship until the age of 7 weeks, with only 6,6% of larvae alive on the age of 8 weeks (negative control). In this way, due to these extreme results in controls (0% of mortality in positive control and 100% of mortality in negative control), the mortality data were also analyzed only between treatments (*I. paraguariensis* and *I. theezans*).

The treatment (*I. paraguariensis* and *I. theezans*), week (1 to 8) and interaction factor (week:treatment) was significantly different among *A. aegypti* larvae mortality for both GLMs models (With and Without positive and negative controls; Table 1). The higher *A. aegypti* larvae mortality was found in Bti treatment (negative control), followed to *I. theezans, I. paraguariensis* and positive control (Table 1; Fig 1a). Also, *I. theezans* hydroalcoholic leaves extract, independently of the extract age, significantly killed more *A. aegypti* larvae them the aqueous *I. paraguariensis* fruits extract (Table 1; Fig 1b).

**Table 1:**
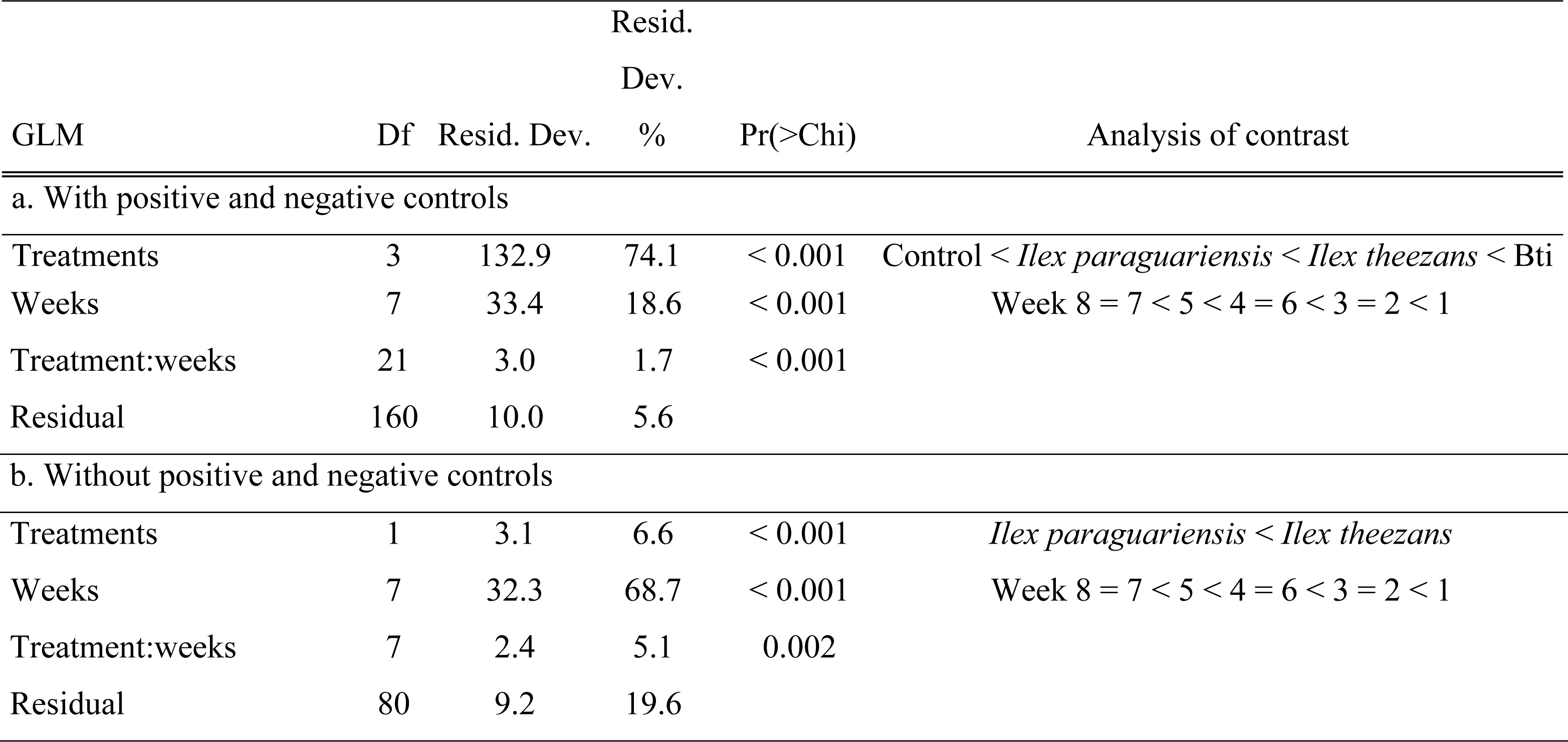
Generalized linear models (GLM), degrees of freedom (Df), Residual Deviance (total and in) and p valuers, comparing the percentage of *Aedes aegypti* larvae mortality after the exposure to treatments (water control, *Bacillus thuringiensis israelensis -*Bti, hydroalcoholic dried leaves extract of *Ilex theezans* and aqueous *Ilex paraguariensis* fruits extract), time (8 weeks) and interaction among treatments and weeks, under laboratory conditions.

**Fig 1.**
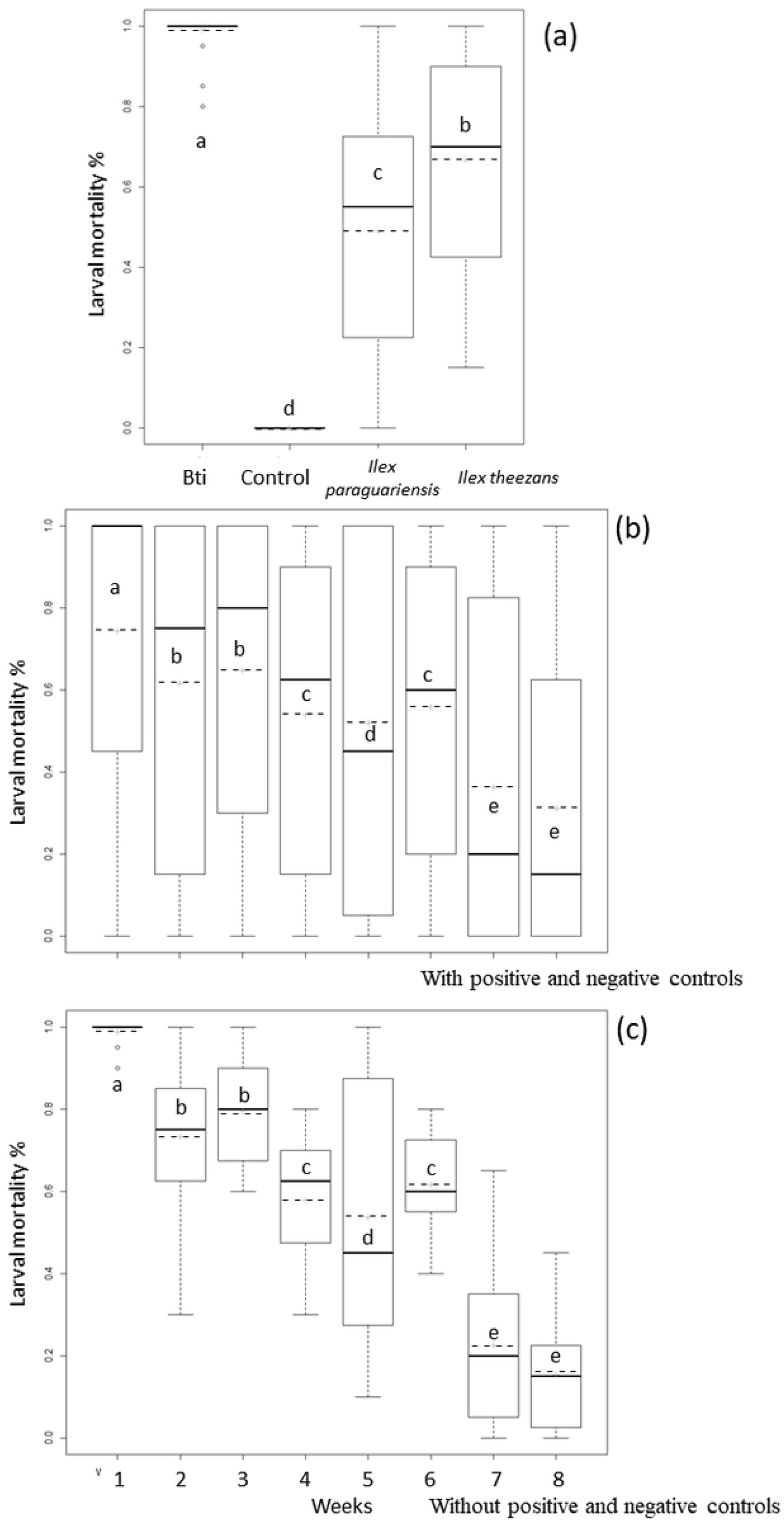
*Aedes aegypti* larvae mortality among treatments (a), sample weeks with positive and negative controls (b) and sample weeks without positive and negative controls (c) among. Different letters (“a”, “b”, “c”, “d” and “e”) indicate significant differences. Boxes represent the quartiles, the bold line represents the median, horizontal dashed line the mean, the vertical dashed line represents the upper and lower limits and circles the outliers.

We observed a positive relationship between *A. aegypti* larvae survivorship and plant extract age. We can also say that in general, both *I. paraguariensis* and *I. theezans* kill less (mainly after 7 weeks) mosquito larvae as the plant extracts age (Fig MS1). We found higher significant in *A. aegypti* larvae mortality in 1 week followed to 2 and 3 weeks, 4 and 6 weeks, 5 weeks and 7 and 8 weeks (Figs 1b and c).

The residual deviance (estimate to explain the variance of the tested variables) in GLM with positive and negative controls, showed that differences in all treatments (74%) was the main responsible for the *A. aegypti* larvae mortality followed by weeks (18%; Table 1). On the other hand, the residual deviance in GLM without positive and negative controls, showed that differences in all weeks (68%) was the main responsible for the *A. aegypti* larvae mortality followed by treatments (19%), and only later was explained by treatments (6%; Table 1).

**Fig MS1:**
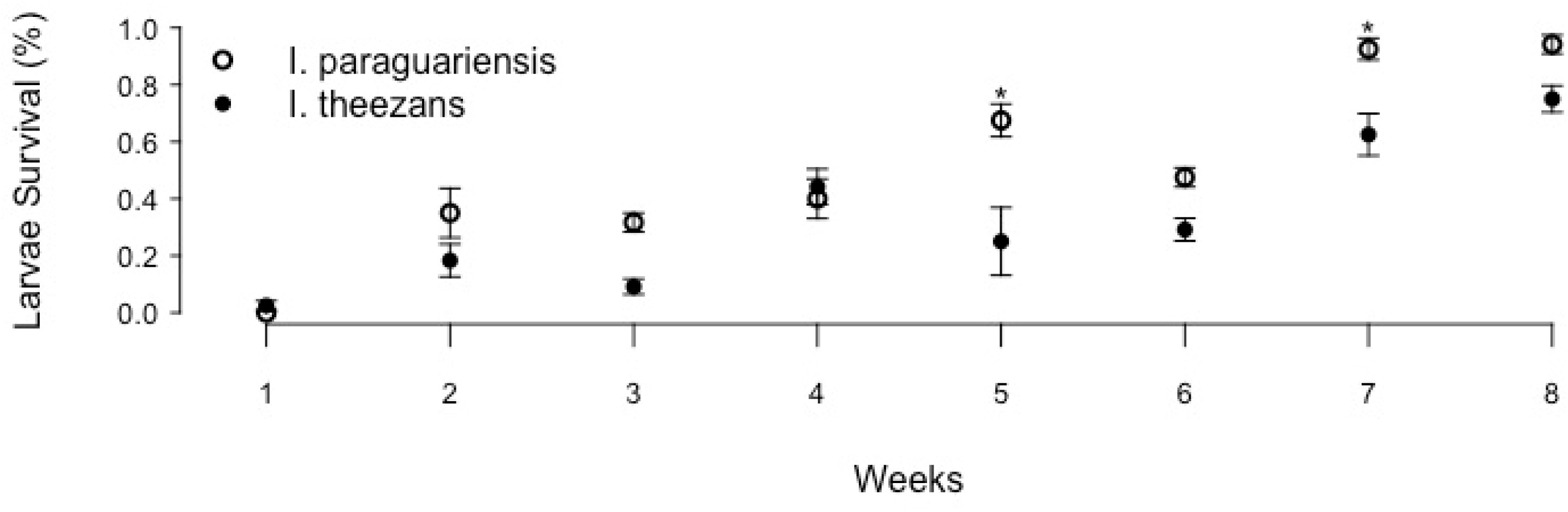
*A. aegypti* larvae survival mean as a function of plant extracts age. The extracts age are represented by weeks. The circles represent, aqueous *I. paraguariensis* fruits extract (open) and *I. theezans* hydroalcoholic leaves extract (closed). The asterisks (*) represents significant statistical difference between *I. paraguariensis* and *I. theezans* affecting larvae survival.

## Discussion

### Reversal of larvicide effect into food resource over time

*I. theezans* and *I. paraguariensis* extracts are promising against mosquito larvae (if applied and monitored in the first weeks) and the possibility of using these plants extracts as larvicides for *A. aegypti* represents. As it is natural extracts and does not leave toxic waste in the environment, its use may be advantageous. This extracts are an abundant and accessible alternative in southern Brazil, where *A. aegypti* infestation and dengue cases have been observed in the last decade [9]. However, extracts age caution should be applied on mosquito control attempts. The main idea of this study is not discouraging the use of such alternative method. We are only concerned in bringing an advice that we need to be aware about the *Ilex* spp. plant and fruit extract age before using it for mosquito control purposes. Better results can also be obtained with the development of additional studies, evaluating the larvicidal activity of pure compounds isolated from these plants. Also, better results can also be obtained as well as evaluate if there is a supporting effect of more than one active principle with larvicidal action for *A. aegypti*.

It is possible that the plant extracts degrade as the time goes on and these organic compounds with previous larvicidal activity may become food for *A. aegypti* larvae (corroborate to increase of importance to residual deviance percentage in GLMs models). The transformation of larvicide effect into food, especially on those treatments seven and eight weeks old, which displayed the highest survivorship rate. Because of this apparently plant extract larvicidal effect loss. Therefore, we recommend that the plant extract age should be considered into attention, caution when using these extracts in time series, since they may serve as food for bacteria [35]. Finally, plant extract age will serve as food for the larvae, contributing to its development rather than combat it, in a negative effect to to mosquito population control.

### A. aegypti larvae mortality between plant extracts

The plant extracts preparation age plays an important role on mosquito larvae mortality (mainly with positive and negative controls). Despite the plant species and parts and extraction method, mosquito larval mortality decrease as the plant extracts age. However, the extracts tested were high efficient in the first weeks of the environment (high mortality). The higher *A. aegypti* larvae mortality found on *I. theezans* extracts can be in partially explained by the use of solvents during the extraction process [36]. The hydroalcoholic extraction method, used for *I. theezans*, extracts reduced polarity chemical constituents from the plant tissues, and these molecules have a greater ability to penetrate the mosquito larvae cells and modify metabolic activities.

On the other hand, aqueous extraction, used for the *I. paraguariensis* fruits, preferentially removes high polarity chemical compounds which are not able to easily penetrate such cells [36]. Also, *A. aegypti* larvae susceptibility to *I. theezans* may be explained by the presence of secondary metabolites of the coumarin class and absence of alkaloids when compared to *I. paraguariensis* [37]. Coumarins are part of several plants secondary metabolism being well known for displaying insecticidal activities, acting as an adult repellent, oviposition deterrence, feeding and growth inhibition, morphogenetic and hormonal system alterations, sexual behavior changes, adult sterilization, among others [38]. It is important to point out that in raw plant extracts the active constituents are usually found in small concentrations [22].

### A. aegypti larvae mortality among week old

In both, *I. theezans* and *I. paraguariensis*, the *A. aegypti* larvae mortality exposed to one-week old plant extracts was reduced to 100%. However, the plant extracts for both plants’ species decrease the mosquito larvae lethality as the plant extracts age. Those results pointed out the need to carefully select the right age for a *Ilex* spp. plant extract before using it for mosquito larvae control purpose. Especially because, regarding to the residual larvicidal power, a product is considered efficient for a pest population control when it reaches a population reduction above 80% of the individuals, otherwise there is the selection of resistance genes [39].

In addition, it is possible that beyond decaying the lethal chemical compounds, which are toxic for mosquito larvae on the first week. As the time goes on the organic compounds present on the plant extracts act as a profitable food source for mosquito larvae. In this way, since organic compounds are well known as being an important component of several larval habitats, forming the basis of many food webs[40,41]. Microorganisms, such as bacteria, play an important role in the cycling and breaking of large organic molecules [42]. Therefore, microorganisms may making them more easily assimilable to aquatic organisms such as mosquito larvae, especially those belonging to the Culicidae family [40]. As a result of this, decomposing microbial communities present a relevant contribution to the diet of culicid larvae, which end up ingested with the organic remains over time [40,43–45].

Our Bti results lead to 93.4% mortality using the eighth week aged solution. The residual effect described in the manufacturer’s technical manual is 30 days. Also, with the values obtained in this work being much higher than those described, with a lethality of 100% for 49 days and 6.6% of larval survival up to 56 days. In this way, it was possible to evaluate the effectiveness of the products that are already being used by the state programs to *A. aegypti* control and combat. Therefore, testing their effectiveness against the existing vector populations, since the information obtained could indicate if there is a need insecticides currently used. It should be emphasized here that the use of the methodology without water renewal in the experiment ends up resulting in a longer residual effect, not evaluating the impact of water renewal in reducing the residual effect [46]. It would be important in other studies to simulate the situation in the field of realities where deposits dominated by permanent emptying and replacement of water, which would probably contribute to reduction of the residual effect duration for all the treatments tested.

## Conclusions

Our study reinforces the idea of the continuously need to implement alternative methods to vector control, but there are long term risk. Also, identifying potential alternative pathways for mosquito population control using natural products originated from the native flora in the short term. The above described plants have high larvicidal potential against the mosquito *A. aegypti*, allowing it’s easy access by the local population, and contributing to the maintenance of the quality of life and well-being of the population. As well as reducing public expenditures with vector control and in the treatment of confirmed cases of dengue. Finally, we found that time would positively affect *A. aegypti* larvae survival due to the plant extracts lethal compounds decay, corroborating our first hypothesis. We also found that the mortality by *B. thuringiensis* var. *israelensis* will be constant throughout the experimental period and *A. aegypti* larvae survival will be lower in the treatments of plant extracts compared to *B. thuringiensis* var. *israelensis.* The high mortality was observed in *I. theezans* compared to *I. paraguariensis*, corroborating our second hypothesis. The strongest residual effect of *I. theezans* was probably due to the presence of chemical on its leaves, such coumarins, hemolytic saponins and cyanogenic glucosides, absent in *I. paraguariensis*.

## Acknowledgements

The authors thank Unochapecó for the availability of the laboratories use and for the Research Program for the Unified Health System.

